# Sox6: A new modulator of renin expression during physiological conditions

**DOI:** 10.1101/556118

**Authors:** Mohammad Saleem, Conrad P. Hodgkinson, Ela W. Contreras, Liang Xiao, Juan A. Gimenez-Bastida, Jason Foss, Alan J. Payne, Maria Mirotsou, Vivian Gama, Victor J. Dzau, Jose A. Gomez

## Abstract

Juxtaglomerular (JG) cells, major sources of renin, differentiate from metanephric mesenchymal cells which give rise to JG cells or a subset of smooth muscle cells of the renal afferent arteriole. During periods of dehydration and salt deprivation JG cells undergo expansion. Gene expression profiling comparing resident renal Mesenchymal Stromal Cells (MSCs) with JG cells indicate that the transcription factor Sox6 is highly expressed in JG cells in the adult kidney. *In vitro*, loss of Sox6 expression reduces differentiation of renal MSCs to renin producing cells. *In vivo*, Sox6 expression is up-regulated during JG cell expansion. Importantly, knockout of Sox6 in Ren1d+ cells halts the increase in renin expressing cells normally seen during JG cell expansion as well as the typical increase in renin. These results support a previously undefined role for Sox6 in renin expression during normal and pathophysiological conditions.

## INTRODUCTION

The renin angiotensin aldosterone system (RAAS) plays a pivotal role in renal development, the control of blood pressure and fluid homeostasis. Inappropriate activation of the RAAS in adult mammals contributes to the development of hypertension, cardiac hypertrophy, and heart failure. The first and rate-limiting step in the initiation of RAAS is catalyzed by the aspartyl protease Renin, which is predominantly produced in the kidney. The mechanisms underlying renin production have been investigated in the developing kidney as well as during pathological conditions in adult animals. In the developing kidney, lineage tracing studies using the Ren1 promoter have shown that renin producing cells derive from mesenchymal precursors at embryonic day (E)11.5 (12, 22, 28). These renin precursors are positive for the transcription factor Foxd1 (12, 19, 26). The expression of renin in the embryo is detectable by E14.5 (17), and increases thereafter, such that by embryonic day 15.5 renin is present in the renal artery, interlobar arteries, and the arcuate arteries of the metanephric kidney (23). After birth, renin expressing cells become restricted to the terminal portion of the juxtaglomerular (JG) afferent arteriole. In adults, conditions such as chronic ischemia, prolonged adrenergic activation, and sodium depletion perturb blood pressure homeostasis and increase the number of renin expressing cells along the afferent arteriole, the kidney interstitium, and inside the glomerulus recapitulating the embryonic distribution of renin expression (9-11). This process of JG cell recruitment occurs via several cellular mechanisms(15, 20, 27). One mechanism involves dedifferentiation and re-expression of renin by renal smooth muscle cells (SMC) along the arteriole (15, 27). An alternative mechanism is the differentiation of pericytes (3, 23, 26) and adult renal mesenchymal stromal cells (MSC) into renin expressing cells (29).

The mechanisms regulating renin expression remain elusive. Various studies have stressed the importance of transcriptional mechanisms in the control of renin expression. Renal smooth muscle cells along the afferent arteriole produce renin and participate in JG cell expansion, and in these cells the transcription factor RBP-J plays a key role in renin expression and the JG cell identity (12). Moreover, transcription factors such as CRAB/CREM, HOX/PBX, and LXRα bind to the renin promoter where they exert positive and negative control over renin expression *in vitro* (4, 14). Also, a recently published study show that super-enhancers are involved to maintain renin-expressing cell identity and memory to preserve multi-system homeostasis (22).

Here we show evidence that the transcription factor Sox6 critically modulates renin expression both *in vitro* and *in vivo*. Sox6, which belongs to the Sry (sex determining region Y) subfamily Sox D proteins (5, 13, 18), activates or represses gene transcription through association with multiple transcription factors (18). Gene centric arrays and genome wide association studies have shown a strong association of Sox6 with hypertension (7, 8, 16, 21). Sox6 is expressed and has biding site in the renin promoter within the super-enhancer region (22). *In vitro*, we found that Sox6 is necessary for the c-AMP-mediated induction of renin expression in renal tissue specific stem-progenitor or mesenchymal stromal cells. In adult mice, JG cell expansion stimulated by a sodium restricted diet and furosemide induces Sox6 expression. Sox6 co-localizes with renin cells; as well as renal MSCs and smooth muscle cells in the afferent arteriole. To validate a role for Sox6 in JG cell expansion/renin expression, we used a novel transgenic mouse model in which Sox6 is specifically ablated in renin expressing cells (Ren1d^Cre^/Sox6^fl/fl^). Utilizing this model, we found that ablation of Sox6 in renin expressing cells halts the increase in renin during JG cell expansion. Taken together, our findings demonstrate a new role for Sox6 in the control of renin expression and suggest a role for Sox6 in hypertension.

## MATERIALS AND METHODS

### Animals

Male C57BL/6 wild type mice eight weeks old (Charles River Laboratories), C57BL/6 Ren1c^YFP^ and Ren1d^Cre^ mice and Sox6^fl/fl^ were used. All animal procedures were approved by Vanderbilt University’s Institutional Animal Care and Use Committee, and the mice were housed and cared for in accordance with the Guide for the Care and Use of Laboratory Animals, US Department of Health and Human Services.

### Transcript profiling

Renal MSCs were isolated from adult C57BL/6 Ren1c YFP mice described above. Juxtaglomerular cells were isolated from adult C57BL/6 Ren1c YFP mice by FACS using YFP expression as a surrogate for renin expression.

### Isolation and culture of renal mesenchymal stromal cells (MSCs)

Renal MSC isolation and culture were carried out as previously reported^14^. Renal MSCs were differentiated with 3-Isobutyl-1-methylxanthine (IBMX −100 µM) and Forskolin (10 µM) (I&F) as described previously ^14,25^.

### Induction of JG cell expansion in vivo

JG cell expansion in vivo was induced by low salt diet plus furosemide as described before ^14^.

### Kidney Immunohistochemistry (IHC)

IHC staining of kidney sections was performed as previously described^28^.

### Flow cytometry

Single cell suspension obtained as before^28^.

### Renin and Sox6 for flow cytometry

Antibodies were labeled with Alexa Fluor antibody labeling kit according to manufacturer’s recommendations. Renin was labeled with Alexa Fluor 488 (Invitrogen cat # A20181), and Sox6 labeled with Alexa Fluor 647 (Invitrogen cat # A20186).

### Western blotting

Kidney tissues were minced with a razor and homogenized with tissue tearor (model # 985370, BioSpec products Inc.) following manufacturer’s instructions. Western blotting was performed as previously described(25).

### In situ hybridization

Localization of Sox6 transcription factor and renin mRNA synthesis was studied using the RNAscope^®^ 2.5 HD duplex detection assay (Advanced Cell diagnostics, ACD, Newark, CA, USA) for in situ hybridization technology following manufacture’s protocols.

### Statistics

All statistical analysis was performed using GraphPad Prism 7.03. A t-test was performed for experiments containing two conditions. One-way or two-way ANOVA was used for experiments with three or more conditions followed by Bonferroni post-hoc tests for comparisons between individual groups. A P-value equal or less than 0.05 was considered significant.

## RESULTS

### Identification of Juxtaglomerular-specific transcription factors

To identify specific transcription factors that are highly expressed in juxtaglomerular (JG) cells, we performed a microarray screen comparing renal progenitor cells (mesenchymal stromal cells (MSCs) and juxtaglomerular cells. As expected, renin and other JG cells markers were highly expressed in JG cells and absent from the MSCs (data not shown). Array results showed that 5573 genes were differentially expressed in JG cells compared to renal CD44+ MSC: 1223 transcripts were up-regulated (20%) and 4350 (80%) down-regulated. Of these 5573 genes, 643 transcripts were involved in transcription, 73 were up-regulated and 570 were down-regulated. Genes up-regulated more than 5 folds are shown (Figure 1a). Sox6 was chosen for its function in controlling cell fate in other systems (2, 13, 18). Microarray fluorescence intensity data (in Arbitrary Units-AU) from MSCs and JG cells samples were plotted. We found that Sox6 was highly expressed in JG cells compared to renal MSCs (Figure 1b). Sox6 microarray data was validated by qRT-PCR (Figure 1c).

**Fig. 1.**
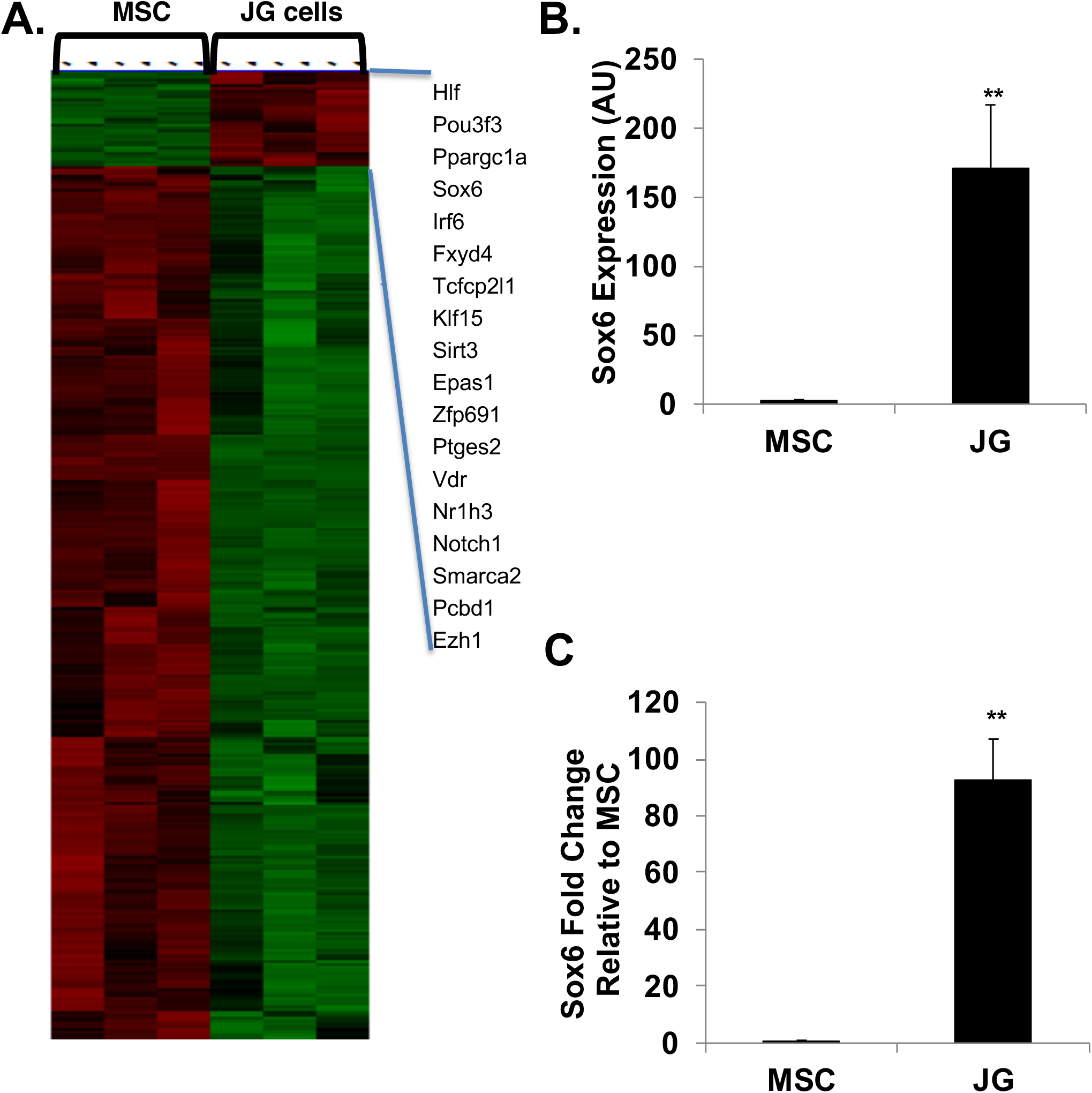
Sox6 expression is up-regulated in JG cells compared to renal MSCs. (**A**) Heat map showing the microarray data comparison between renal CD44+ MSCs and JG cells. Data analyzed using d-Chip, which focuses on transcriptional regulators. Genes upregulated ≥ 5-fold in JG-cells are presented in right side of heat map. (**B**) Microarray expression data of Sox6 in renal MSCs and JG cells. (**C**) Sox6 qRT-PCR validation of microarray data. Expression values for Sox6 are shown relative to GAPDH. At least four independent samples per group. P values calculated using a student unpaired t-test. **P<0.01.

### Sox6 is involved in renin expression in vitro

Our gene expression array data indicated that Sox6 is highly expressed in cells that produce renin. To validate this in a model that induces renin expression, CD44+ tissue specific MSCs were isolated from C57BL6 Ren1c YFP adult kidneys and treated with 3-Isobutyl-1-methylxanthine (IBMX − 100 µM) and Forskolin (10 µM) (I&F) to differentiate into renin expressing cells as previously described (29). This treatment concomitantly increased both renin and Sox6 expression (Figure 2a and b), suggesting that Sox6 may control renin expression. We investigated this possibility by reducing Sox6 levels with a specific shRNA in renal MSCs isolated from Ren1c YFP adult mice (Figure 2c and d). Control non-targeted shRNA and cAMP Responsive Element Modulator (CREM) specific shRNA were used as a negative and positive controls respectively. Knockdown of Sox6 prevented MSC differentiation into renin producing cells in response to IBMX and forskolin to same levels as the positive control CREM (Figure 2d). This indicates that Sox6 is necessary for renin expression in renal MSCs induced by cAMP.

**Fig. 2.**
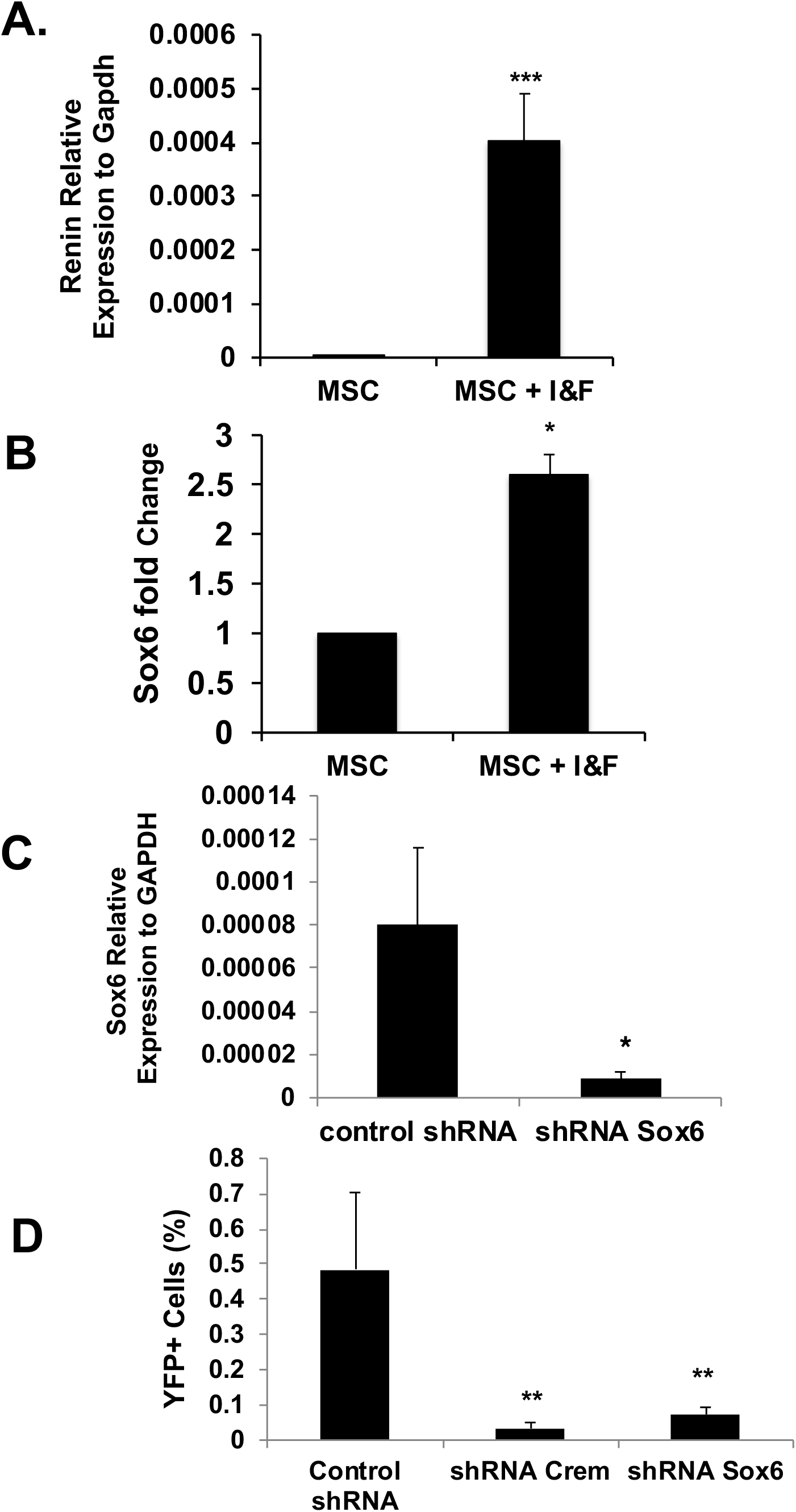
Sox6 is essential for cAMP induction of renin expression *in vitro*. Renal CD44^+^ Mesenchymal Stromal Cells (MSC) were isolated from adult wild-type mice. Quantitative RT-PCR analysis of renin and Sox6 expression after IBMX and forskolin (I&F) treatment was performed after 7 days of treatment. N= 4, Data are presented as the mean ± SEM. P calculated with an unpaired t-test. *P< 0.05, *** P< 0.001. (**A**) Renin. (**B**) Sox6. Renal CD44^+^ Mesenchymal Stromal Cells (MSC) were isolated from C57BL6 Ren1c YFP adult mice and cultured in growth medium for three to five passages before use, and YFP expression used as surrogate for renin expression. (**C**) Quantitative RT-PCR analysis of Sox6 knockdown in MSC, three independent experiments. Data are presented as the mean ± SEM. P calculated with an unpaired t-test. *P< 0.05. (**D**) Down regulation of Sox6 affects the differentiation of renal MSC to renin expressing cells. Three independent experiments, data are presented as the mean ± SEM. P value calculated using an unpaired t-test. **P< 0.01 comparing Sox6 shRNA to control shRNA or Crem shRNA to control shRNA.

### Sox6 expression increases during juxtaglomerular cell expansion

To unravel the role of Sox6 in the kidney and its contribution to regulation of blood pressure homeostasis, we initially analyzed Sox6 expression in adult mice. We found that Sox6 is expressed in the cortex and medulla of the adult kidney (Figures 3a and b). Specificity of the Sox6 antibody was demonstrated with tissue from Sox6 knockout mice (2)and a specific peptide to discard false positive staining (Supplementary Figure S1A). After LowNa/Fu treatment, Sox6 was co-localized with renin expressing cells in the glomeruli and afferent arteriole (Figures 3c and d). Co-localization of Sox6 staining was also observed with cells involved in JG recruitment such as vascular and perivascular MSCs (24) as well as alpha smooth muscle actin positive (αSMA+) smooth muscle cells (SMCs) in the afferent arteriole (28) (Supplementary Figure S1B and C). Confirming our previous study, renin expression was significantly up-regulated following feeding a low sodium diet and treatment with furosemide (LowNa/Fu), which also induces juxtaglomerular (JG) cell expansion (29) (Figure 3e). Concomitant with 3-fold increase in the number of renin expressing cells (renin+ cells), Sox6 expressing cell (Sox6+ cells) increased 36-fold (Figure 3e and f). To further support a role for Sox6 in controlling renin expression, we performed chromatin immunoprecipitation (ChIP) assay on freshly isolated kidney cells following LowNa/Fu treatment. From *in silico* analysis of the renin promoter we found four putative Sox6 binding sites within −5Kb from transcription start site. Analysis of this region by ChIP indicated that Sox6 was bound to the renin promoter in renal cells during JG cell expansion (Supplementary Figure S2). These results support the notion that Sox6 modulates renin expression during JG cell expansion; controlling renin expression during this process.

**Fig. 3.**
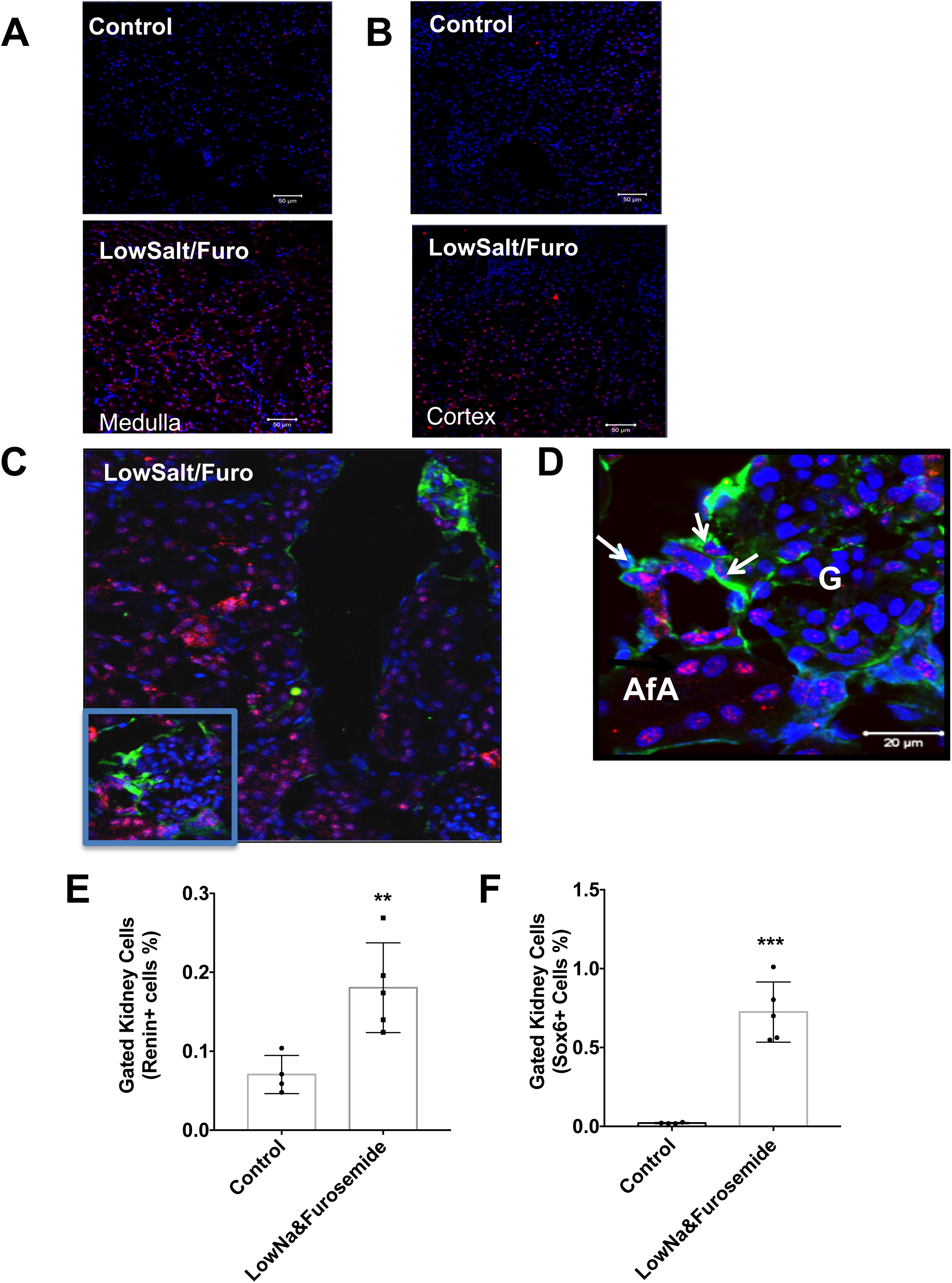
Sox6 expression is up regulated during JG cell expansion. C57BL/6 wild type (6-8 weeks old) were administered a Low Sodium diet (0.02% NaCl) plus furosemide, (drinking water −2.28 mmol/L) for ten days. Control animals received normal chow (0.6% NaCl). Representative 20X confocal images of kidney medulla and cortex sections stained with Sox6 (red) and nucleus (blue) from control mice. Upper panels: control untreated mice; lower panels: LowNa/Furo treated mice. Scale bar 50 µm. Representative 20X confocal image of kidney sections stained with Sox6 (red), and nucleus (blue) (**A**) Kidney medulla and (B) Kidney cortex. (C) Representative 20X confocal images of kidney sections stained with Sox6 (red), Renin (green), and nucleus (blue) from low sodium diet and furosemide treated mice. (**D**) Magnified image of the glomeruli showing the co-expression of renin and Sox6. Arrows point to Sox6+ Renin+ cells. G: glomerulus; AfA: afferent arteriole. Scale bar 20 µm. Flow cytometry analysis of isolated mouse kidney cells using specific antibodies to renin and Sox6 to quantify renal cells expressing these proteins. (**E**) Renin expression in mice before and after low sodium and furosemide treatment. (**F**) Sox6 expression in mice before and after low sodium and furosemide treatment. N= 4 to 5, Data are presented as the mean ± SEM. P calculated with an unpaired t-test. **P< 0.01, *** P< 0.001 low-salt/furosemide versus no treatment.

### Renin expression in low salt diet and furosemide (LowNa/Fu), and non-treated animals

To further define a role for Sox6 in controlling renin expression during physiological conditions, we developed a new transgenic mouse by crossing the Ren1d^Cre^ transgenic mouse (27) with the Sox6^fl/fl^ mouse (6) (Figure 4a and supplementary figure S3). The resulting Ren1d^Cre^/Sox6^fl/fl^ mice lack Sox6 specifically in renin expressing cells. To elaborate a role of Sox6 in renin expression we measured renin protein by Western blot after LowNa/Fu treatment. In the LowNa/Fu treated group, levels of renin protein expression were significantly higher in wild-type (Ren1d^cre^/Sox6^wt/wt^) compared to Sox6 specific knockout (Ren1d^cre^/Sox6^fl/fl^) mice (Figure 4b). Quantitatively, renin expression in the Ren1d^cre^/Sox6^fl/fl^ did not increase after stimulation with sodium restricted diet and furosemide; moreover, renin expression was significantly lower compared to the wild-type mice. In non-treated group, levels of renin expression were decreased about 72% in Sox6 specific knock out compared to wild-type mice. Within the wild-type groups, there was significant difference in the levels of renin expression in non-treated compared to LowNa/Fu treated mice (Figure 4b and c). As another parameter of JG cell expansion, we measured plasma renin activity after LowNa/Fu treatment. Mice with specific ablation of Sox6 in renin expressing cells had a significantly lower plasma renin activity compared to wild-type mice (Figures 4d). To define if Sox6 affects renin transcription, RNA expression levels were measured by in situ hybridization using specific probes to renin and Sox6 mRNA. At baseline the number of renin and Sox6 double positive cells in the juxtaglomerular area is significantly lower in Ren1d^Cre^/Sox6^fl/fl^ mice compared to Ren1d^Cre^/Sox6^wt/wt^ mice (Figure 4e-f and Supplementary Figure 5sA and C). Following low sodium and furosemide treatment renin mRNA levels and in situ signal intensity increased in the Ren1d^Cre^/Sox6^wt/wt^ mice, and the number of glomeruli with renin and Sox6 double positive cells increase (Figure 4e and f). However, there was no increase in renin mRNA levels in the Ren1d^Cre^/Sox6^fl/fl^ mice as well as Sox6 and renin double positive cells (Figure 4e-f and Supplementary Figure 5sB and D). The above results indicate that Sox6 plays a role in renin expression control; influencing both mRNA and protein levels.

**Fig. 4.**
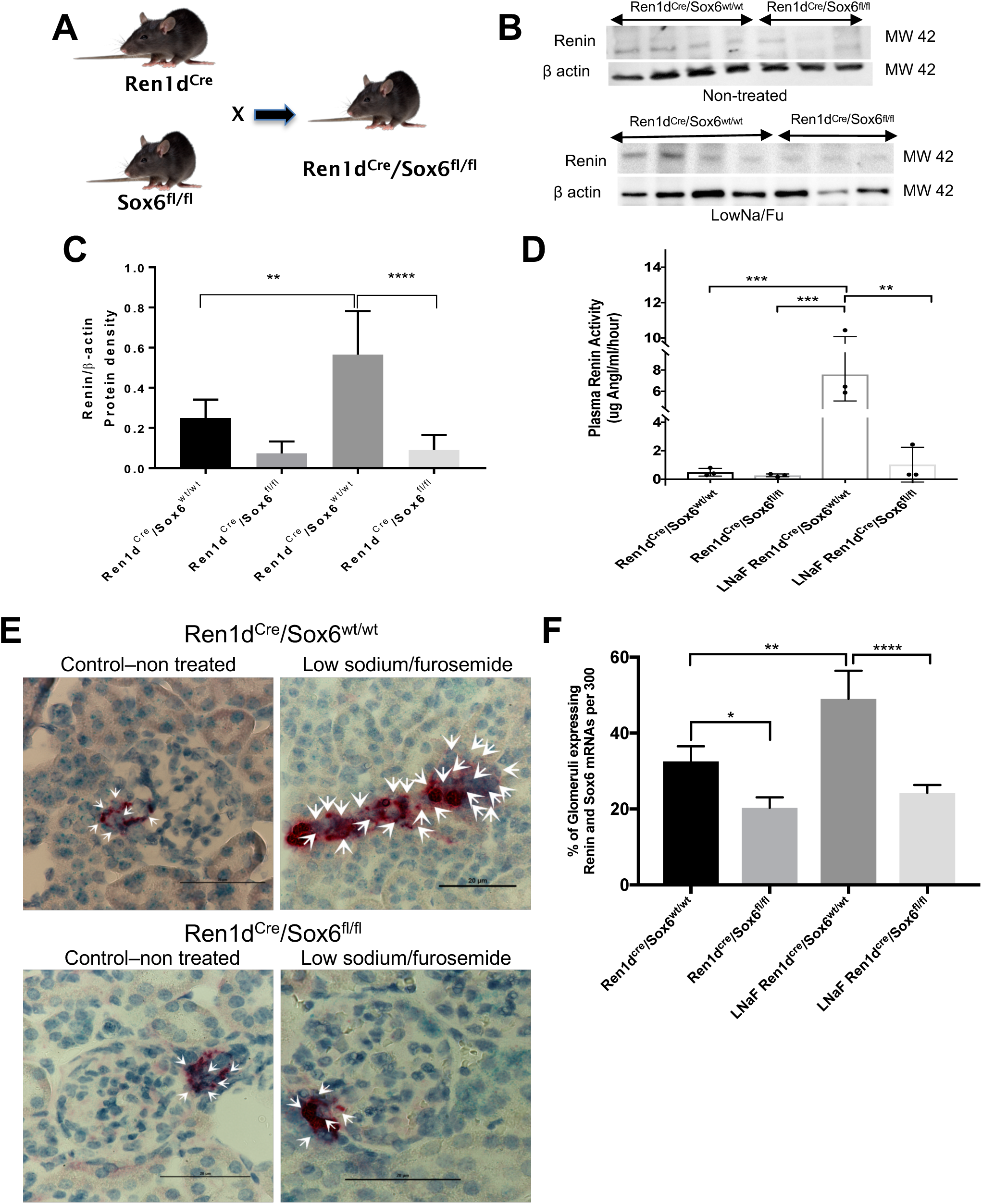
Specific knockout of Sox6 in JG cells inhibits the increase in renin expression during JG cell expansion. (**A**) Schematic representation of development of Ren1d^Cre^/Sox6^fl/fl^. (B) After ten days of lowNa/Fu treatment, kidneys were isolated and Western blot was performed. Representative blots showing levels of renin expression in non-treated (upper blot) and lowNa/Fu treated (lower blot). β-actin was used as a loading control. (**C**) Quantification of immunoblots. N= 5 to 7, Data are presented as the mean ± SEM. P calculated with one-way ANOVA followed by Tukey post-hoc test. **P< 0.01, **** P< 0.0001. (**D**) Plasma was isolated from non-treated and lowNa/Fu treated Ren1d^Cre^/Sox6^fl/fl^ and Ren1d^Cre^/Sox6^wt/wt^ mice (6-8 weeks old) mice, and plasma renin activity (PRA) determine as μg Ang I/mL/h. Data are presented as mean ± SEM. N=3 per group, and P calculated with one-way ANOVA followed by turkey post-hoc test. **P< 0.01, *** P< 0.001. (**E**) Specific knock out of Sox6 in JG cells inhibits the increase in renin mRNA expression cells during JG cell expansion. Ren1d^Cre^/Sox6^fl/fl^ and Ren1d^Cre^/Sox6^wt/wt^ (6-8 weeks old) were administered a Low Sodium diet (0.02% NaCl) plus furosemide ((LowNa/Fu) drinking water 2.28 mmol/L) for ten days. After treatment, kidneys were isolated and perfusion-fixed with 10% neutral buffered formalin solution, dehydrated in a graduated ethanol series, and embedded in paraffin. Renin and Sox6 mRNAs amplification were performed by following manufacturer’s instructions. Red and green punctuated dots represent renin and Sox6 mRNAs expression respectively. Representative microscopy images. 60X magnification. Scale bar 20 microns. Arrows point at cells expressing both renin and Sox6 mRNAs. (**F**) In situ hybridization quantification of glomeruli expressing both renin and Sox6 mRNAs from a total at least three hundred glomeruli per sample. Specific probes were used to detect renin and Sox6. N= 4, Data are presented as the mean values ± SEM. P was calculated with one-way ANOVA followed by Tukey post-hoc test. * P<0.05, ** P<0.01, **** P<0.0001.

### Specific knock out of Sox6 in renin expressing cells prevents increase in renin expression during JG cell expansion

Next, we measured the number of renin expressing cells *in vivo* during conditions that promote JG cell expansion using flow cytometry. When compared to control mice, the Ren1d^Cre^/ Sox6^fl/fl^ mice failed to increase the number of renin expressing cells in response to a sodium deficient diet and furosemide (Figure 5b to e). CD44 and CD73 were used as markers of renal MSC or renal stem progenitor cells. We found that there was an increase in the number of renin and CD44 double positive cells and a decrease in the number of renin and CD73 positive cells in mice lacking Sox6 in renin-expressing cells (Supplementary Figure S5B and C). This suggests that ablation of Sox6 *in vivo* affects differentiation of MSC to renin expressing cells and may affect the maintenance of the progenitor phenotype after differentiation to renin expressing cells (Supplementary Figure S5B and C). Furthermore, our results showed that Sox6 is important for renin expression by smooth muscle cells during JG cell expansion (Figure 5e, Supplementary Figure S5A). Expression of Sox6 is up-regulated after changes in renin expression induced by LowNa/Fu treatment (Figure 3f). Specific ablation of Sox6 in renin expressing cells inhibited the increase in renin producing cells during JG cell expansion (Figure 4c), as well as the number of Sox6+ and renin+ double positive cells (Figure 5d). Taken together, the results presented here allow us to conclude that Sox6 has a new function in renin expression control.

**Fig. 5.**
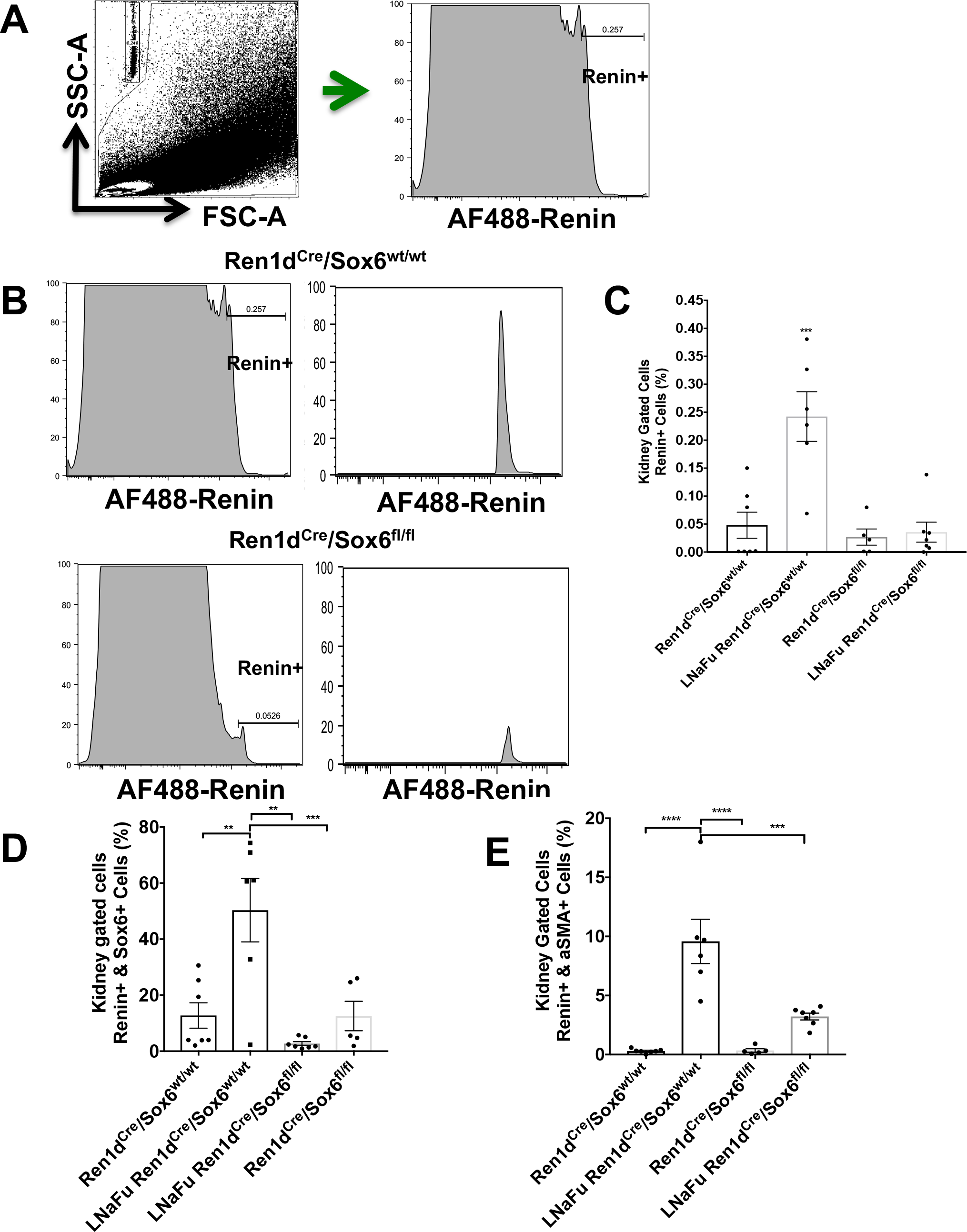
Specific knock out of Sox6 in JG cells inhibits the increase in renin expressing cells during JG cell expansion. Ren1d^Cre^/Sox6^fl/fl^ and Ren1d^Cre^/Sox6^wt/wt^ (6-8 weeks old) were administered a Low Sodium diet (0.02% NaCl) plus furosemide (LowNa/Fu), (drinking water −2.28 mmol/L) for ten days. Flow cytometry analysis of isolated mouse kidney cells using specific antibodies to renin and Sox6 to quantify renal cells expressing these proteins. (**A**) Left panel, representative flow cytometry dot plots of kidneys from Ren1d^Cre^/Sox6^fl/fl^ and Ren1d^Cre^/Sox6^wt/wt^ showing gating strategy. Right panel, representative flow cytometry histograms of kidneys from Ren1d^Cre^/Sox6^fl/fl^ and Ren1d^Cre^/Sox6^wt/wt^ showing gating for renin expressing cells (Alexa Fluor 488–renin+). **(B)** Renin expressing cells histograms in kidneys after lowNa/Fu treatment, upper panels Ren1d^Cre^/Sox6^wt/wt^ mice and lower panels Ren1d^Cre^/Sox6^fl/fl^ mice. Bar graphs showing kidney cells positive for the expression of (**C**) Renin, (**D**) Renin and Sox6 double positive cells, and (**E**) renin and alpha-SMA double positive cells. N= 6 to 7. Data are presented as the mean ± SEM. P calculated with an unpaired t-test. *P< 0.05, **P< 0.01 ***P < 0.001.

## DISCUSSION

Renin is the rate-limiting enzyme to produce ang II, and thus plays a pivotal role in the RAAS system. RAAS activity is essential for water and electrolytes balance and blood pressure control. RAAS exerts its blood pressure control through plasma circulating renin. While renin is produced in different tissues, the predominant site of production is the kidney, where it is produced and stored by Juxtaglomerular cells (1). Control of renin expression and secretion is of great importance for human health. Renin expression and secretion is carefully coordinated by several proteins that work in a harmonized manner maintaining blood pressure and fluid homeostasis.

In this study, we demonstrate that Sox6 has a new function in the control of renin expression. Under pathological conditions, JG cells arise from several cellular sources. Previously, we reported that resident renal mesenchymal stromal cells (MSC) play a role in JG cell expansion and that a subset of these MSCs differentiates into renin expressing cells (29). Other studies have stressed the importance of arteriolar smooth muscle cells (SMCs) in afferent arteriole (9, 22, 27). In our study, we show that not only does Sox6 co-localize with renin, but it also co-localizes with markers of both renal MSCs and SMCs during JG cell expansion. Likewise, flow cytometry quantification of renin positive cells showed that Sox6 specific ablation in renin expressing cells inhibits the increase in renin+ cells during JG cell expansion. Moreover, we found that Sox6 controls the expression of renin in MSCs after cAMP stimulus *in vitro*.

Gene centric array and genome-wide association (GWAS) studies have identified an association between hypertension in various ethnic groups and single nucleotide polymorphisms (SNPs) in the Sox6 gene (7, 8, 16, 21). It is currently unknown how these SNPs affect Sox6 function. In the light of our study, it is possible that these SNPs affect the ability of Sox6 control over renin expression.

The second messenger cAMP is important for renin expression. The renin promoter contains cAMP responsive elements in the proximal and distal promoters where cAMP responsive transcription factors bind promoting renin expression (4). In silico analysis of renin promoter shows that there is a Sox6 binding site in the proximal promoter, and not within any of the cAMP responsive elements. However, Sox6 expression itself is stimulated by cAMP and its promoter contains binding sites for cAMP responsive transcription factor such as ATF1, ATF2, and ATF7 which may bind and stimulate Sox6 expression. Future experiments defining the role of cAMP in Sox6 expression and its relationship to renin expression will be of interest.

*In vivo*, renin mRNA expression increased after 10 days of low sodium diet and furosemide. Sox6 specific ablation in renin expressing cells inhibited the increase in renin protein expression and mRNA expression observed in wild type mice during JG cell expansion.

Taken together, our findings indicate that Sox6 has a previously undefined role in modulating renin expression at baseline and in response to sodium and volume deprivation. Given this critical function, Sox6 might also be a therapeutic target for the treatment of hypertension.

## ACKNOWLEDGEMENTS

We would like to thank Dr. R. Ariel Gomez for kindly providing us with the Ren1dCre mice, and Dr. Monique Lefebvre from the Cleveland Clinic for generously providing the Sox6^fl/fl^ transgenic mice. We would like to thank Dr. David G. Harrison for his comments to the manuscript.

## DISCLOSURES

No conflicts of interest, financial or otherwise, are declared by the authors.

### Author Contribution

J.A.G., and M.S. conceived and designed the work presented in this manuscript. J.A.G and M.S executed most experiments and analyzed most of the data. J.A.G drafted, revised and approved the final version of the manuscript. C.P.H., E.W.C-P., L.X., J.A.G-B., J.F., and A.J.P., provided technical expertise. M.M. and J.A.G. prepared and analyzed microarray data. M.S., and J.A.G. wrote the manuscript. M.S., V.G., C.P.H., and V.J.D. commented on the manuscript and provided key technical support.

