## Supplemental data for "Sox6: A new modulator of renin expression during physiological conditions" (from another version of this preprint: version 1, 2019-02-20)

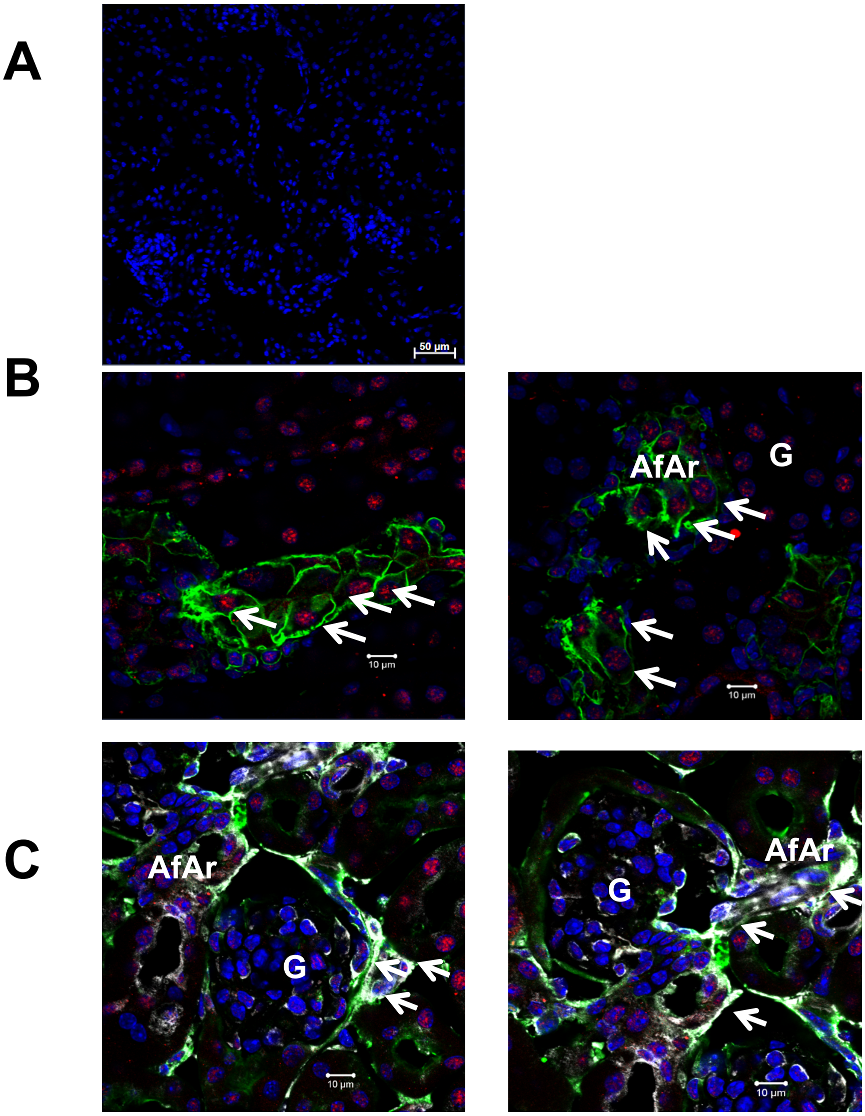


**Supplementary Figure S1.** C57BL/6 wild type (6-8 weeks old) were administered a Low Sodium diet (0.02% NaCl) plus furosemide, (in drinking water -2.28 mmol/L). (A) Representative 20X confocal image of kidney section stained with Sox6 antibody and Sox6 peptide-competition assay. Sox6 peptide was used as control to determine specificity of the antibody in the kidney sections. (B) Representative 63X confocal image of kidney sections stained with Sox6 (red), CD44 (green), and nucleus (blue) from LowSalt/Furo treated mice. Arrows point to Sox6+ CD44+ cells. G: glomerulus; AfA: afferent arteriole. Scale bar 10 µm. (C) Representative 63X confocal image of kidney sections stained with Sox6 (red), αSMA (cyan), and nucleus (blue) from control non treated (left panels) and LowSalt/Furo treated (right panels) mice. Arrows point to Sox6+ αSMA + cells. G: glomerulus; AfA: afferent arteriole. Scale bar 10 µm. Confocal microscopy analysis four mice per condition with three 10 µm kidney sections per kidney and multiple confocal images per section.

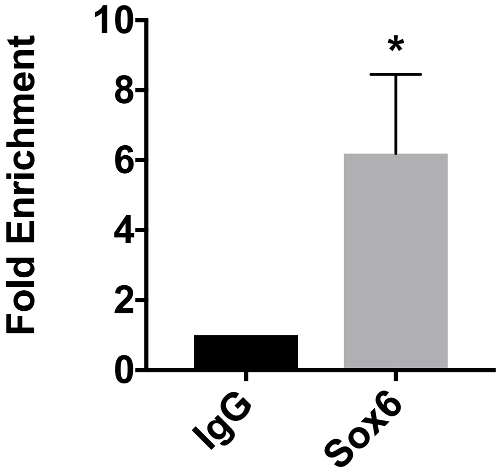


**Supplementary Figure S2.** Chromatin immunoprecipitation ChIP. ChIP assays were performed according the manufacturer's instructions (Cell Signaling, SimpleChIP Enzymatic Chromatin IP kit #9003). Mice kidneys were digested with collagenase as described above, and single cells suspension nuclei were digested with 0.1ul Micrococcal nuclease per $4x10^6$ cells (amount of Micrococcal nuclease was empirically determined according the manufacturer's instructions). Immunoprecipitation was performed with ChIP validated antibodies: rabbit IgG control (Cell Signaling, #2729) and Sox6 (ThermoFisher #PA5-34616). Immunoprecipitated DNA was quantified by qPCR (ThermoFisher, Power SYBR Green PCR Master Mix, #4367659) with primers for the Renin promoter (Qiagen, EpiTect ChIP qPCR Primer Assay for Mouse Ren1, NM_031192.2 (-)01Kb): catalogue number GPM1028245(-)01A). A serial dilution of the 2% input chromatin DNA (undiluted, 1:5, 1:25, 1:125) was used to create a standard curve and determine efficiency of amplification. Percent input was calculated and negative control IgG values subtracted. Data is presented as the fold change of percent input between IgG and Sox6 samples. C57BL/6 wild type (6-8 weeks old) were administered a Low Sodium diet plus furosemide, (LowNa/Furo) for 10 days. After treatment kidneys were isolated, collagenase treated to obtain single cells isolation. Cells were lysed and the nuclear fraction assayed for Sox6 binding to renin promoter by chromatin immunoprecipitation using a specific Sox6 antibody. Binding of Sox6 to renin promoter ChIP presented as fold enrichment. N=3. Data are presented as the mean ± SEM. P calculated with an unpaired t-test. *P< 0.05.

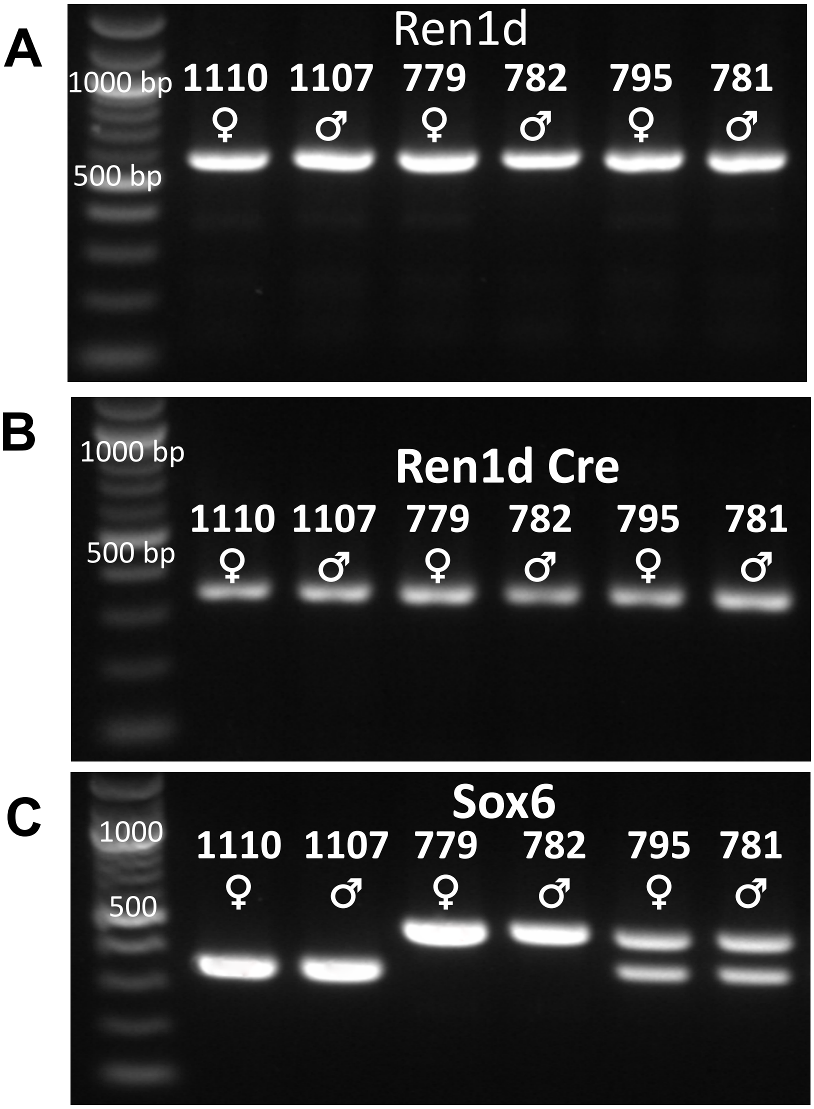


**Supplementary Figure S3.** Tail tips were cut using alcohol-sterilized scissors and put in an Eppendorf tude. Then, 500 µL of solution 2 (DNA extraction solution II Cat # 200600-14-Stratagene) was added to the tail tip with 2 µL pronase K per sample (cat # 200600-82) and incubated at 55 $^{0}$C overnight. Next morning, tubes were spin down for 30 s on bench top centrifuge and chilled on ice for 10 min. After adding and mixing 200 µL of solution 3, tubes were incubated on ice for 5 min and were centrifuged at 20000 g at 4 $^{0}$C to get pellet. DNA was precipitated by adding 2 volumes of 100% ethanol to the pellet and centrifuged at 20000 g at 4 $^{0}$C for 10 min. Lastly, water (100 µL) was added in obtained pellet with DNA that was used for PCR reactions for genotyping. Ren1d$^{cre}$ and Sox6$^{Fl/Fl}$ mice were crossed to get Ren1d$^{cre}$Sox6$^{Fl/Fl}$ transgenic mouse. **A**. Ren1d runs in 1.5% agarose gel at size 585 bp. **B**. Ren1d$^{cre}$ runs in 1.5% agarose gel at size 341 bp. **C.** Sox6 runs in 1.5% agarose gel at size 450 bp (Sox6 fl/fl, one band) 450 and 360 bp (Sox6 fl/-, two bands) and 363 bp (Sox6 wild-type ).

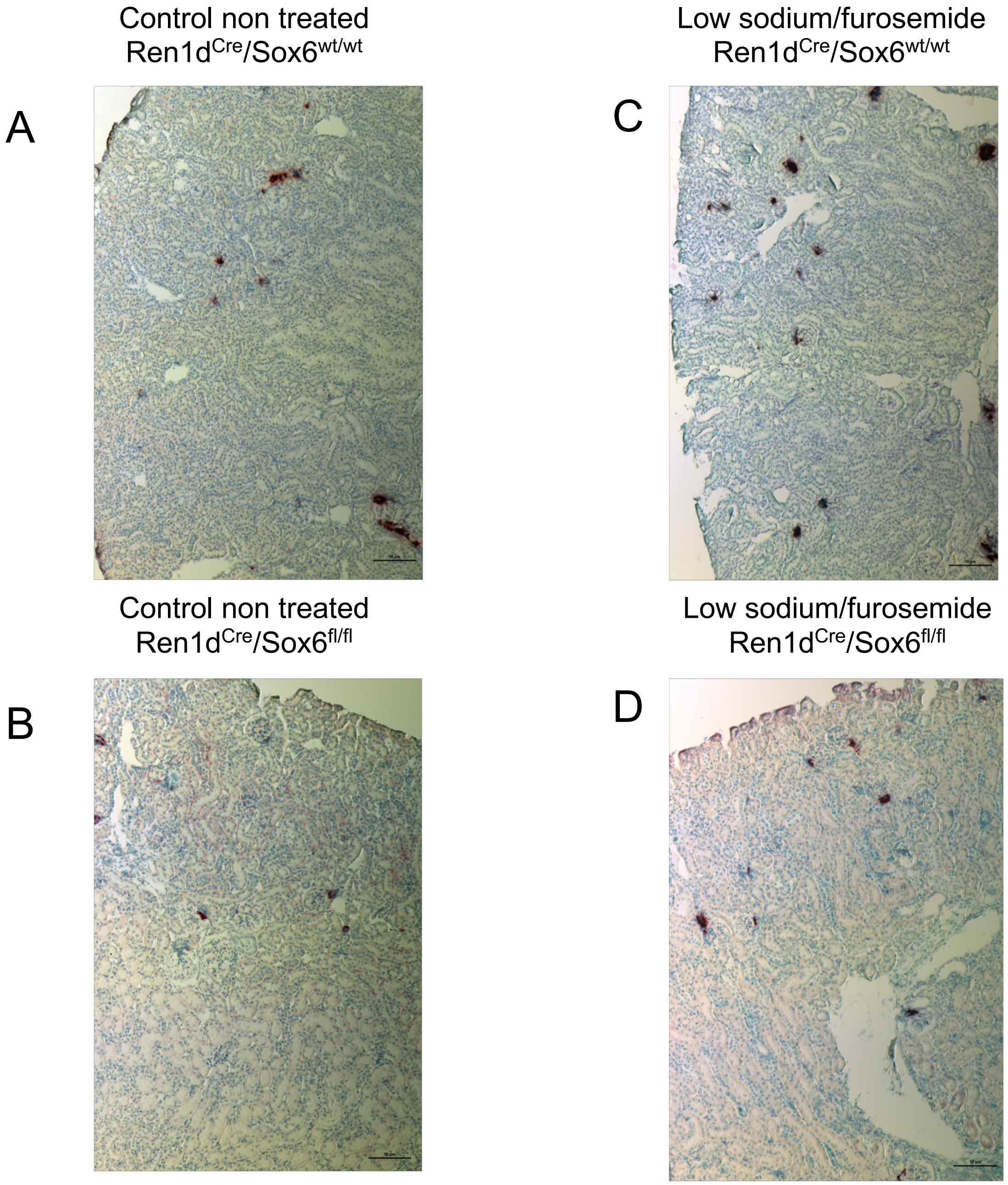


**Supplementalry Figure S5.** Specific knock out of Sox6 in JG cells inhibits the increase in renin mRNA expression cells during JG cell expansion. Ren1d$^{Cre}$/Sox6$^{fl/fl}$ and Ren1d$^{Cre}$/Sox6$^{wt/wt}$ (6-8 weeks old) were administered a Low Sodium diet (0.02% NaCl) plus furosemide ((LowNa/Fu) drinking water - 2.28 mmol/L) for ten days. After treatment, kidneys were isolated and perfusion-fixed with 10% neutral buffered formalin solution, dehydrated in a graduated ethanol series, and embedded in paraffin (For detailed description, see method section). Renin and Sox6 mRNAs amplification was performed by following manufacture's instructions. Red and green punctuated dots representing renin and Sox6 mRNAs expressions respectively. Representative microscopy images. 10X magnification. Scale bar 50 microns.

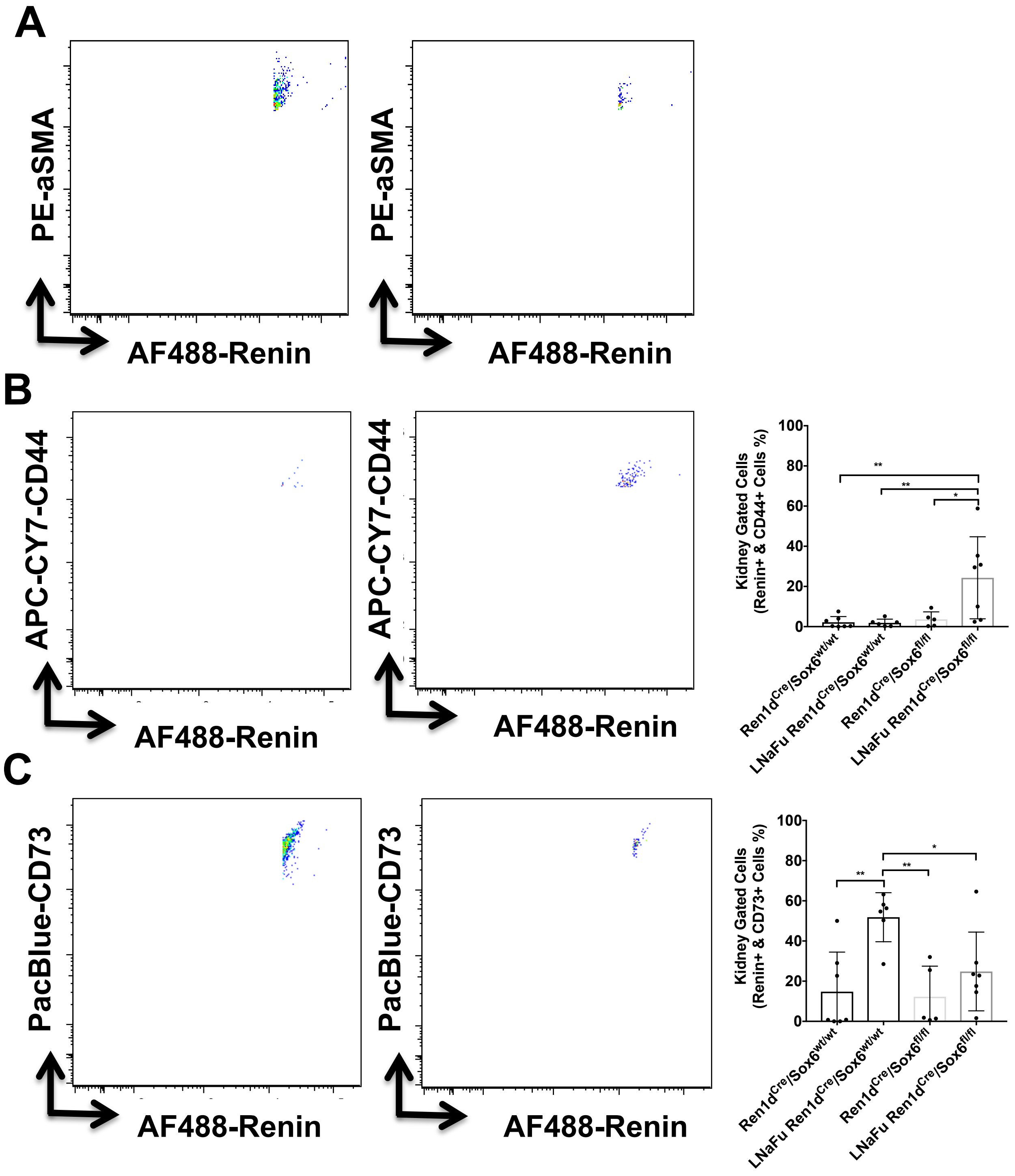
A
PE-aSMA
AF488-Renin
PE-aSMA
AF488-Renin
B
APC-CY7-CD44
AF488-Renin
APC-CY7-CD44
AF488-Renin
Kidney Gated Cells
(Renin+ & CD44+ Cells %)
100
80
60
40
20
0
**
**
*
Ren1dCre/Sox6wt/wt
LNaFu Ren1dCre/Sox6wt/wt
Ren1dCre/Sox6fl/fl
LNaFu Ren1dCre/Sox6fl/fl
C
PacBlue-CD73
AF488-Renin
PacBlue-CD73
AF488-Renin
Kidney Gated Cells
(Renin+ & CD73+ Cells %)
100
80
60
40
20
0
*
**
**
Ren1dCre/Sox6wt/wt
LNaFu Ren1dCre/Sox6wt/wt
Ren1dCre/Sox6fl/fl
LNaFu Ren1dCre/Sox6fl/fl

**Supplementary Figure S5.** Specific knock out of Sox6 in JG cells inhibits the increase in renin expressing cells during JG cell expansion. Ren1d$^{Cre}$/Sox6$^{fl/fl}$ and Ren1d$^{Cre}$/Sox6$^{wt/wt}$ (6-8 weeks old) were administered a Low Sodium diet (0.02% NaCl) plus furosemide, (drinking water -2.28 mmol/L) for ten days. Flow cytometry analysis of isolated mouse kidney cells using specific labeled antibodies to AlexaFluor488-renin, AlexaFluor647-Sox6, APC-Cy7-CD44, PacificBlue-CD73 and PE-alpha smooth muscle actin (aSMA) to quantify renal cells expressing these proteins. CD44 and CD73 were used as markers for renal mesenchymal stem cells. After treatment, kidneys were harvested for flow cytometry. (A) Left panel bar graph showing double positive renin and aSMA expressing cells in mice after low sodium and furosemide treatment. Middle and right panels flow cytometry dot plots showing renin and aSMA double positive cell population. (B) Left panel bar graph showing double positive renin and CD44 expressing cells in mice after low sodium and furosemide treatment. Middle and right panels flow cytometry dot plots showing renin and CD44 double positive cell population. (C) Left panel bar graph showing double positive renin and CD73 expressing cells in mice after low sodium and furosemide treatment. Middle and right panels flow cytometry dot plots showing renin and CD73 double positive cell population. N= 6 to 7. Data are presented as the mean ± SEM. P calculated with an unpaired t-test. *P< 0.05, **P < 0.01.

SUPPLEMENTARY METHODS

*Transcript profiling*. Renal MSCs were isolated from adult C57BL/6 Ren1c YFP mice described above. Juxtaglomerular cells were isolated from adult C57BL/6 Ren1c YFP mice by FACS using YFP expression as a surrogate for renin expression. Total RNA from the purified MSCs and JG cell population was isolated using a PicoPure Arcturus kit (Invitrogen) according to the manufacturer’s instructions. Complimentary cDNA was generated with an Ovation Pico WTA System V2 kit (NuGEN) that maintains stoichiometry of the original RNA population. The Affymetrix Mouse 430 2.0 array was used for gene profiling. Data analysis focused on transcriptional and epigenetic regulators and performed by using Dchip, David or Toppgene. Filtering was done by the following criteria: a) Variation across samples: 1 < standard deviation/mean < 1000; and b) P call % in the arrays used >= 20%. Agglomerative hierarchical clustering of MSCs and JG cell samples was done with Partek Genomics Suite 6.6 (Partek Inc., St Louis, MO, USA) using the Euclidian similarity measure and average linkage as the linkage method. Probes that showed the highest difference in mean expression compared to MSCs baseline in any of the JG samples was chosen, and the expression validated using qRT-PCR. The raw and processed data was uploaded to the NIH GEO database and has been assigned the Accession number GSE114903.

*Sox6 shRNA knock down*. Renal MSCs were infected with virus encoding specific shRNA to Sox6, and CREM or a non-targeting control shRNA (Mission shRNA from Sigma). After infection, cells were selected with puromycin for 3 days and cultured until confluence. I&F treatment as described above. After I&F treatment for 7 days, cells were fixed with paraformaldehyde, and the expression of YFP was measured and quantified using Cellomics ArrayScan, and iDev software (ThermoFisher Scientific) respectively. False negatives were discarded based on image analysis on the combination of Dapi (nuclei) and YFP expression channels.

*Kidney Immunohistochemistry (IHC).* IHC staining of kidney sections was performed as previously described[28]. The following primary antibodies were used: anti-CD44 (1/50 dilution, 103001, BioLegend); anti-renin (1/100 dilution, kindly provided by Dr. Tadashi Inagami, Vanderbilt University); anti-renin (1/50 dilution, 1206, Innovative Research); anti-αSMA (1/500 dilution, A5228, Sigma); and anti-Sox6 (1/500, ab30455, abcam) [24]. Secondary antibodies were Alexa 594 goat anti-rabbit IgG (A- 11012); Alexa 488 donkey anti-goat IgG (A-11055); Alexa 647 donkey anti-mouse IgG (A-31571); and Alexa 488 goat anti-rat IgG (A-11006), all secondary antibodies were from Invitrogen. Nuclei were counterstained with DAPI concentration (1 ug/mL). Sox6 peptide (ab30530-abcam) was used as blocking peptide during IHC to determine the specificity of the Sox6 antibody (ab30530). Immunofluorescence staining sections were visualized and photographed using Zeiss 510 Inverted Confocal Microscope or Axiovert 200 microscope.

*Flow cytometry*. Single cell suspension obtained as before[28], and stained with renin Alexa fluor 488 labeled antibody (R&D Systems, AF4277), Sox6 Alexa fluor 647 (abcam, ab30455), aSMA PE labeled (marker of renal arteriolar smooth muscle cells), CD44 APC-Cy7 (marker of renal mesenchymal stromal cells) and/or CD73 Pacific-blue (marker of renal mesenchymal stromal cells) for 1hr at 4oC in FACS buffer (1xPBS, 0.5% w/v BSA, 2mM EDTA). After incubation with antibodies, Cells washed with FACS buffer and analyzed on a FACSCanto II flow cytometer. Gating was based on flow minus one and single stained sample controls. Flow cytometry data was analyzed with FlowJo version 10.

*Renin and Sox6 antibodies for flow cytometry*. Antibodies were labeled with Alexa Fluor antibody labeling kit according to manufacturer's recommendations. Renin was labeled with Alexa Fluor 488 (Invitrogen cat # A20181), and Sox6 labeled with Alexa Fluor 647 (Invitrogen cat # A20186).

*Western blotting*: Kidney tissues were minced with a razor and homogenized with tissue tearor (model # 985370, BioSpec products Inc.) following manufacturer's instructions. Western blotting was performed as previously described [28.] Primary antibody dilutions used were 1:500 for Sox6 (#Ab-30455, Abcam, MA, USA) and renin (Santa Cruz Biotechnology, SCBT, USA), while 1: 2500 for β-actin (Sigma, MO, USA). Samples were resolved on 4–15% Tris-glycine gels (Bio-rad, 456-1083, CA USA) by SDS/PAGE. Gels were transblotted onto polyvinylidene fluoride membrane for 2 h at 4 °C. Membranes were then blocked with a mixture of milk (5%) and BSA (3%) in TBST at room temperature (RT) for 1 h and probed with respective primary antibodies in TBST overnight at 4 °C. After washing with TBST, membranes were incubated with corresponding secondary antibodies conjugated with horseradish peroxidase (HRP). Chemiluminescent reagent was used (Bio-rad, 1705060, CA, USA) to visualize protein bands using ChemiDoc™ XRS+ System image station (Bio-rad, 1708265, CA, USA). Protein bands were quantified using software integrated with the image station.

*In situ hybridization*: Localization of Sox6 transcription factor and renin mRNA synthesis was studied using the RNAscope® 2.5 HD duplex detection assay (Advanced Cell diagnostics, ACD, Newark, CA, USA) for in situ hybridization technology following manufacture's protocols. Briefly, kidneys were perfusion-fixed with 10% neutral buffered formalin solution, dehydrated in a graduated ethanol series, and embedded in paraffin. Ten micrometer sections were cut. Sections were de-paraffinized in histo-clear solution at room temperature (RT), followed by blocking of endogenous peroxidase with RNAscope H2O2. Tissue was retrieved by boiling in target retrieval solution at 100-104oC for 15 min, then treated with protease plus at 40oC for 30 min. Target probes (designed by ACD: Mm-Sox6, Ref # 472061-C1; Mm-Ren1, Ref # 433461-C2) were hybridized for 2 h at 40oC, followed by a series of signal amplification (Amp 1-6) and in between washing with RNAscope wash buffer. Red (renin mRNA) and green (Sox6 mRNA) were detected after 6 and 10 signal amplifications respectively. RNA staining signal was identified as red and green punctuate dots. All hybridization steps at 40oC were performed in a HybEZ Hybridization System (ACD). After the completion of RNAscope assay, tissue sections were counterstained with 50% hematoxylin staining solution for 30 s at RT, followed by mounting with VectaMount mounting medium (H-500, Vector laboratories, Burlingame, CA, USA) and drying slides in a dry oven at 60oC for 30 min. The tissue sections were viewed with Nikon Eclipse Ti, (Software NIS-Elements AR 4.40.00 64-bit). Randomization and Blinding: Two separate blinded investigators carried out the staining and counting the glomerulus containing renin and Sox6 double positive cells. Investigators were blinded with respect to animal identifiers and group assignments. Values were averaged between the two blinded investigators.
